# Crystal structure of a human plasma membrane phospholipid flippase

**DOI:** 10.1101/2019.12.23.881698

**Authors:** Hanayo Nakanishi, Katsumasa Irie, Katsumori Segawa, Kazuya Hasegawa, Yoshinori Fujiyoshi, Shigekazu Nagata, Kazuhiro Abe

**Author notes:** Correspondence and requests for materials should be addressed to K.A.

## Abstract

ATP11C, a member of P4-ATPase flippase, exclusively translocates phosphatidylserine from the outer to the inner leaflets of the plasma membrane, and maintains the asymmetric distribution of phosphatidylserine in the living cell. However, the mechanisms by which ATP11C translocates phosphatidylserine remain elusive. Here we show the crystal structures of a human plasma membrane flippase, ATP11C-CDC50A complex, in an outward-open E2P conformation. Two phosphatidylserine molecules are in a conduit that continues from the cell surface to the occlusion site in the middle of the membrane. Mutations in either of the phosphotidylserine binding sites or along the pathway between significantly impairs specific ATPase and transport activities. We propose a model for phosphatidylserine translocation from the outer to the inner leaflet of the plasma membrane.

## Introduction

Phospholipids are asymmetrically distributed between the outer and inner leaflets in the plasma membrane of eukaryotic cells. Aminophospholipids such as phosphatidylserine (PtdSer) and phosphatidylethanolamine (PtdEtn) are confined to the inner leaflet, while phosphatidylcholine and sphingomyelin are enriched in the outer leaflet^1^. The asymmetric distribution of phospholipids is widely conserved in eukaryotes, being tuned for barrier functions and various signal transductions on the plasma membrane^2^. On occasion, this phospholipid asymmetry in the plasma membrane is disrupted, exposing PtdSer on the cell surface. Cells undergoing apoptosis expose PtdSer as an “eat me” signal to phagocytes^3,4^. Activated platelets also display PtdSer as a scaffold for clotting enzyme reactions^5^. The amphipathic nature of phospholipids prevents their spontaneous flip-flop movement across the lipid bilayer in most cases, and the translocation of phospholipids therefore needs membrane proteins to overcome the energetic barrier required for the phospholipid translocation^2^.

While scramblases mediate non-specific and bi-directional movement of phospholipids between inner and outer leaflets^6^, flippases exhibit ATP-driven, directional and up-hill translocation of phospholipids from the outer to inner leaflets against their concentration gradient across the membrane bilayer^7,8,9^. Different from other members of cation-transporting P-type ATPases^10–13^, Type IV P-type ATPase (P4-ATPase) comprises a subfamily of P-type ATPases that transports phospholipid^14^. Among all 14 members of P4-ATPase in humans, ATP11A and ATP11C work as aminophospholipid-specific flippases at the plasma membrane^3,4^. They require an accessary subunit, CDC50A, for their correct localization and the functional expression on the plasma membrane^15–17^. In fact, cells lacking ATP11A and ATP11C, or CDC50A almost completely lose flippase activity for PtdSer and PtdEtn at the plasma membrane, resulting in failure to maintain the asymmetric accumulation of PtdSer in the inner leaflet. Their inactivation causes PtdSer exposure on the cell surface. In apoptotic cells, ATP11A and ATP11C are subjected to a caspase-mediated proteolysis and irreversible inactivation. It is proposed that calcium ions likely also inhibit the flippase activity for calcium-dependent PtdSer exposure in activated platelets or lymphocytes^18^. Regulation of aminophospholipid asymmetry by flippases is physiologically important; ATP11A-deficient mice are embryonic lethal^19^, and ATP11C-deficient mice display pleiotropic phenotypes such as B-cell lymphopenia^20,21^, cholestasis^22^, mild anemia^23^, and dystocia. Recently, mutations in the ATP11C gene were identified in patients suffered from anemia^24^.

The directional translocation of specific phospholipids mediated by P4-ATPases is achieved by the cyclic conversion of enzyme conformations, E1, E2 and their auto-phosphorylated forms E1P and E2P, similar to the Post-Albers type reaction scheme^14,25^ for cation-transporting P-type ATPases. However, despite the huge size of phospholipids relative to inorganic cations, the mechanism by which P4-flippases translocate phospholipids across the membrane, the so-called “giant substrate problem” ^26,27^, has remained elusive and yet is of considerable interest. Here, we describe a crystal structure of a *bona fide* human plasma membrane flippase ATP11C-CDC50A complex in the outward-open E2P conformation, analyzed to 3.9 Å resolution. Our structure has two PtdSer molecules simultaneously bound in the putative lipid translocating conduit. The structure, together with functional analyses, enable us to propose a molecular mechanism for the phospholipid translocation across the plasma membrane by ATP11C.

## Results and Discussion

### Overall structure of the outward-open conformation

Human ATP11C and CDC50A were expressed using the BacMam system (Fig. S1, Methods)^28^. Purified and deglycosylated proteins were mixed with dioleoylphosphatidylcholine (DOPC)^29^, and crystallized in the presence of phosphate analogue^30^ beryllium fluoride (BeF_x_) and dioleoylphosphatidylserine (DOPS). Crystals were harvested in the presence of excess DOPS, which was key to preservation of crystal quality. X-ray diffraction data from more than 1,500 individual crystals were merged, and the structure was determined by molecular replacement using the atomic model of the E2BeF state of ATP8A1^31^ as a search model, at a resolution of 3.9Å with acceptable statistics of R_work_/R_free_ = 27.9/34.7 (Fig. S1, Table S1). As is seen in most of the other crystallized P-type ATPases, molecular packing occurs as type I crystals in which complexes are embedded in the lipid bilayer^32^. The asymmetric unit of the crystal consists of four protomers of ATP11C-CDC50A. Due to different crystal contacts, the appearance of the electron density map differs significantly for each protomer (Fig. S2). Despite limited resolution, however, well-ordered regions, especially the transmembrane (TM) region of protomer A and B were visible at side chain level (Fig. S3). The overall molecular conformations of the four protomers are essentially the same, although structural variations in some of the loop structures exist (Fig. S2). We therefore focus on the well-ordered protomer A in what follows.

Like other members of the well-characterized, cation-transporting P-type ATPases^11–13^, the up-hill translocation of aminophospholipids by ATP11C is achieved according to the Post-Albers type reaction scheme^14,25^ (Fig. 1A). The outward-open E2P conformation captures PtdSer or PtdEtn on the outer leaflet and induces dephosphorylation of E2P, thus PtdSer- or PtdEtn-dependent ATP hydrolysis can be detected (Fig. 1B), similar to the inward transport of K^+^ by Na^+^,K^+^-ATPase and H^+^,K^+^-ATPase. As we included BeF_x_ and PtdSer for the crystallization, the molecular conformation^33^ was expected to be the outward-open E2P state in which PtdSer is bound to the conduit facing to the exoplasmic side (Fig. 1A). In fact, the overall structure of the ATP11C-CDC50A complex (Fig. 1C) is very close to the corresponding E2P structures of recently reported flippases yeast Drs2p-Cdc50p^34^ and human ATP8A1-CDC50A^31^ complexes. The sequence identities of each catalytic subunit are 35.3% and 36.2%, respectively (Fig. S4 for sequence alignment)^35^. The molecular conformation is also defined by the relative orientations of the cytoplasmic domains^36^ (actuator (A), phosphorylation (P) and nucleotide-binding (N) domains), and the arrangement of the ten transmembrane (TM) helices of the catalytic subunit ATP11C. The phosphate analogue BeF_x_ likely forms a covalent bond to the invariant aspartate in the ^409^DKTG signature sequence, which is covered by the ^179^DGES/T motif located on the surface of the A domain to prevent spontaneous dephosphorylation by the bulk water (Fig. S5). In this conformation, the N domain is segregated from the P domain, and is expected to be relatively flexible compared to the other two domains due to the lack of intimate inter-domain interactions, and this is consistent with its poor electron density (Fig. S2,3). The relative orientation of the A and P domains in ATP11C is close to those observed in Drs2p and ATP8A1 E2P states, and clearly different from that in the ATP8A1 E2-P_i_ transition state (Fig. S5), indicating that the present ATP11C structure adopts an E2P state.

**Fig. 1.**
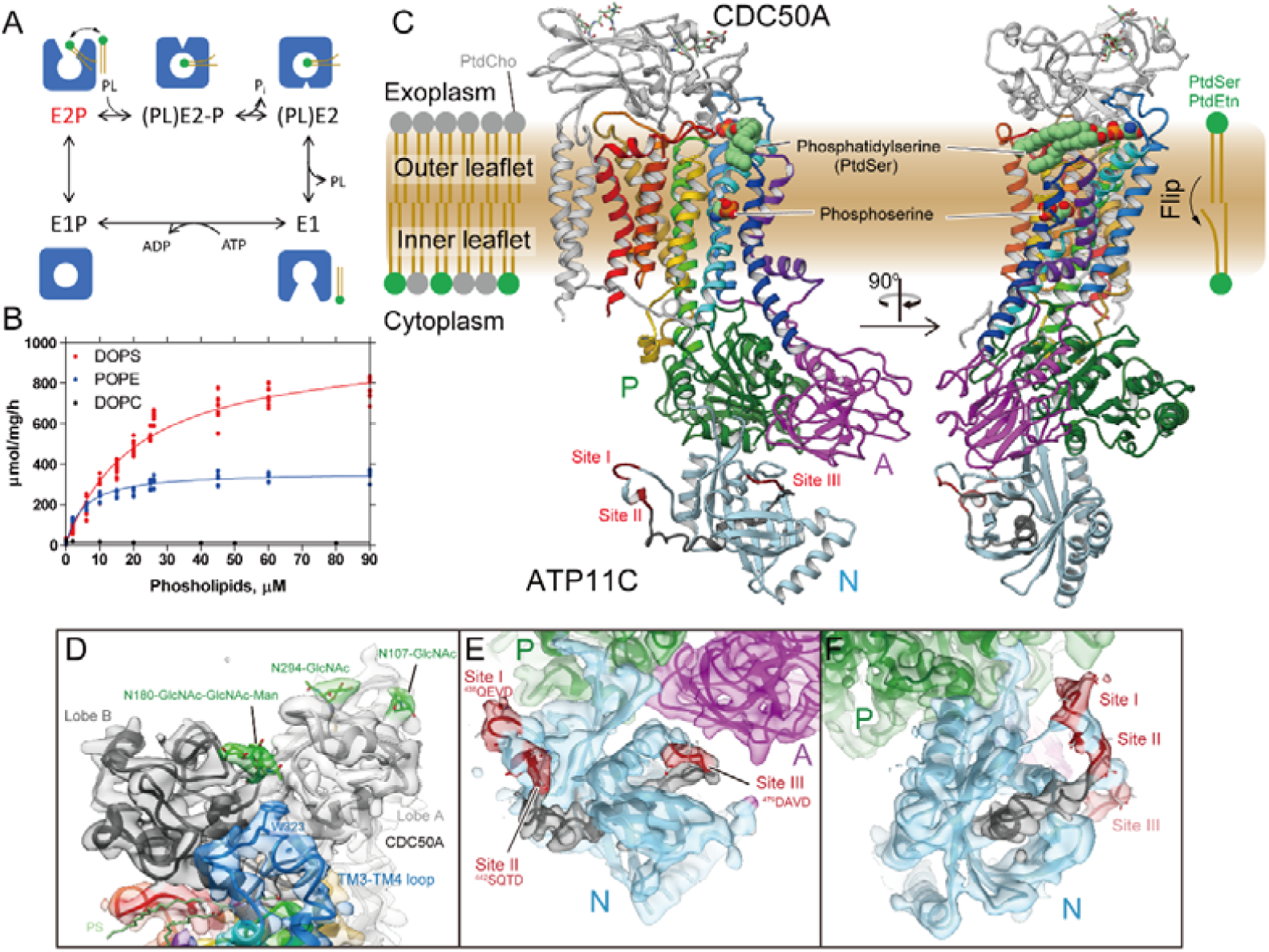
Crystal structure of ATP11C-CDC50A complex. (A) Reaction scheme of phospholipid (PL) translocation coupled with ATP-hydrolysis. Cartoons represent molecular conformations of ATP11C-CDC50A complex (inward- or outward-, and open or occluded states). (B) PL-dependent ATPase activity by the purified ATP11C-CDC50A complex. Specific ATPase activities in the presence of DOPS (red), POPE (blue) or DOPC (black) were plotted as a function of their concentrations. DOPS gave the highest ATPase activity, POPE showed intermediate, while there was no detectable ATPase activity in the presence of DOPC. (C) Overall structure of the outward-open E2P state of ATP11C-CDC50A complex in ribbon representation. Color of the ATP11C gradually changes from the N-terminus (purple) to the C-terminus (red). CDC50A subunit is shown in grey ribbon, and attached *N*-linked glycans were displayed as green sticks. A DOPS molecule (PtdSer) and its hydrophilic phosphoserine part are shown as spheres in the exoplasmic cavity and the occlusion site in the middle of the TM domain, respectively. Three cytoplasmic domains are indicated with different colors, and caspase recognition sequences at the N domain surface are indicated in red. Gray ribbon in the N domain thus indicates the region that is removed after the caspase cleavage. (D) Closed view of the exoplasmic region of CDC50A subunit. Surface represents electron density map (1.5σ). Lobe A (light grey) and lobe B (dark grey) are shown in different colors, and TM3-4 loop is shown in blue with Trp323 side chain fitted into the map. *N*-linked glycans (green) are highlighted. (E,F) Close-up of the N domain (light blue) from two different viewpoints. Surface shows electron density map at 1.0σ contour level. Three caspase recognition sites (I, II and III) and the region in between them (Gly466-Asn477) are indicated in red and grey, respectively.

The CDC50A subunit consists of two TM helices with a long N-terminal tail and a short C-terminus on the cytoplasmic side, and a large exoplasmic domain in which four *N*-linked (Asn107, Asn180, Asn190 and Asn294) and one *O*-linked (Ser292) glycosylation sites are located. We extensively investigated various combinations of glycosylation site mutants to improve protein expression and crystal quality, and found that the Asn190Gln/Ser292Trp double mutant produced the best crystals. Because the samples were treated with endoglycosidase to remove excess glycans during purification, an acetylglucosamine (GlcNAc) moiety is retained on each three remaining *N*-linked glycosylation sites. Interestingly, PNGase treatment, which truncates all glycans including core GlcNAc, produced tiny crystals, suggesting that these GlcNAcs contribute to rather stable crystal packing. We modeled one GlcNAc moiety for each of the two sites (Asn107, Asn294), and three glycans at a position close to Asn180 (Asn180-GlcNAc-GlcNAc-Man). The three glycans at Asn180 fit into a groove formed between two lobes (lobe A and B) of CDC50A and Trp323 (in the connecting loop between TM3 and TM4 of ATP11C and conserved in mammalian flippases that requires CDC50A), like a wedge, thereby keeping them together and sterically protecting them from the endoglycosidase during purification (Fig. 1D).

The electron density map of the N domain allowed visualization of its overall folding at C_α_ level, and made it possible to assign the location of the three caspase recognition sites (Fig. 1E,F). Inactivation of ATP11A or ATP11C by effector caspases requires cleavage of multiple caspase sites located on the N domain, and a single site cleavage is not enough for full inactivation of the flippase^4,18^. There are three caspase-recognition sites in ATP11C (site I – III, Fig. 1, Fig. S4), and these are located on loops at both ends of an α-helix-containing region. Site I and site II are very close to each other at the N-terminal side of the helix, and site III is at the C-terminal side. This region, especially the α-helical part in between caspase recognition sites, is located at the center of the N domain scaffold, and seems like a bolt that holds together the surrounding α-helix and several β-sheets, and thus obviously important for N domain folding. Interaction between this helix and surrounding N domain segments may be sufficient to keep its fold even if one of the caspase sites is cleaved. Cleavage at both ends of the helix must lead to irreversible unfolding of the N domain which is required for the ATP binding. Note, all three sites are exposed on the same side of the N domain with a distance of around 35∼40Å in between two regions (site I,II and site III). Such geometry of cleavage sites suggests that instantaneous two-site cleavage may occur by a caspase-3 dimer^37^ upon apoptotic signal transduction, rather than single, independent cleavages, given that the distance between the catalytic center of the caspase-3 dimer is around 40Å.

### Outward-open conformation

According to the reaction scheme of P4-flippase (Fig. 1A), the outward-open E2P state, mimicked by the BeF_x_-binding, is a reaction state responsible for the incorporation of PtdSer from the outer leaflet to the transport occlusion site. A structural requirement for the outward-open conformation is therefore the transient formation of a conduit that physically allows the translocation of the phospholipid head group from the outer leaflet to the occlusion site in the middle of the membrane. We found a longitudinal crevice in the peripheral region of TM in ATP11C (Fig. 2A, Fig. S6). The crevice is composed of TM2, TM4 and TM6 helices and runs along TM4, and is continuous from the exoplasmic surface of the lipid bilayer to the middle of the membrane (Fig. 4B). Thus, the unwound part of TM4 (^356^PVSM motif), which has been implicated in the lipid transport^27^, is exposed to the hydrophobic bulk lipid. Comparison with other flippase structures in the corresponding reaction state reveals that the crevice in ATP11C is wider than that in Drs2p activated form^34^ (Fig. S6). In the ATP8A1 E2P state, a membrane crevice is not formed at all^31^, probably due to its longer C-terminal regulatory domain compared with that in ATP11C (∼38 amino acids)^38^ (Fig. S4). The exoplasmic side of the crevice is closed in the PtdSer-occluded E2-P_i_ transition state of ATP8A1, and clearly different from that in the ATP11C E2P state (Fig. S6). In ATP11C, TM2 is kinked at Pro94, an amino acid residue conserved in all mammalian P4-ATPases (Fig. 2B, Fig. S6), so that the exoplasmic side of TM2 departs from the central axis of the crevice in this region. This structural feature enables the formation of the wide and continuous crevice from the exoplasmic side to the PtdSer occlusion site near the unwound part of TM4. Substitution of Pro94 for alanine significantly reduces apparent affinities for both PtdSer and PtdEtn (Fig. 2C), indicating that the TM2 kink is key for developing a crevice structure that is wide enough for passage of a phospholipid head group. We conclude that the observed longitudinal crevice is in fact the outward facing conduit that enables the phospholipid translocation.

**Fig. 2.**
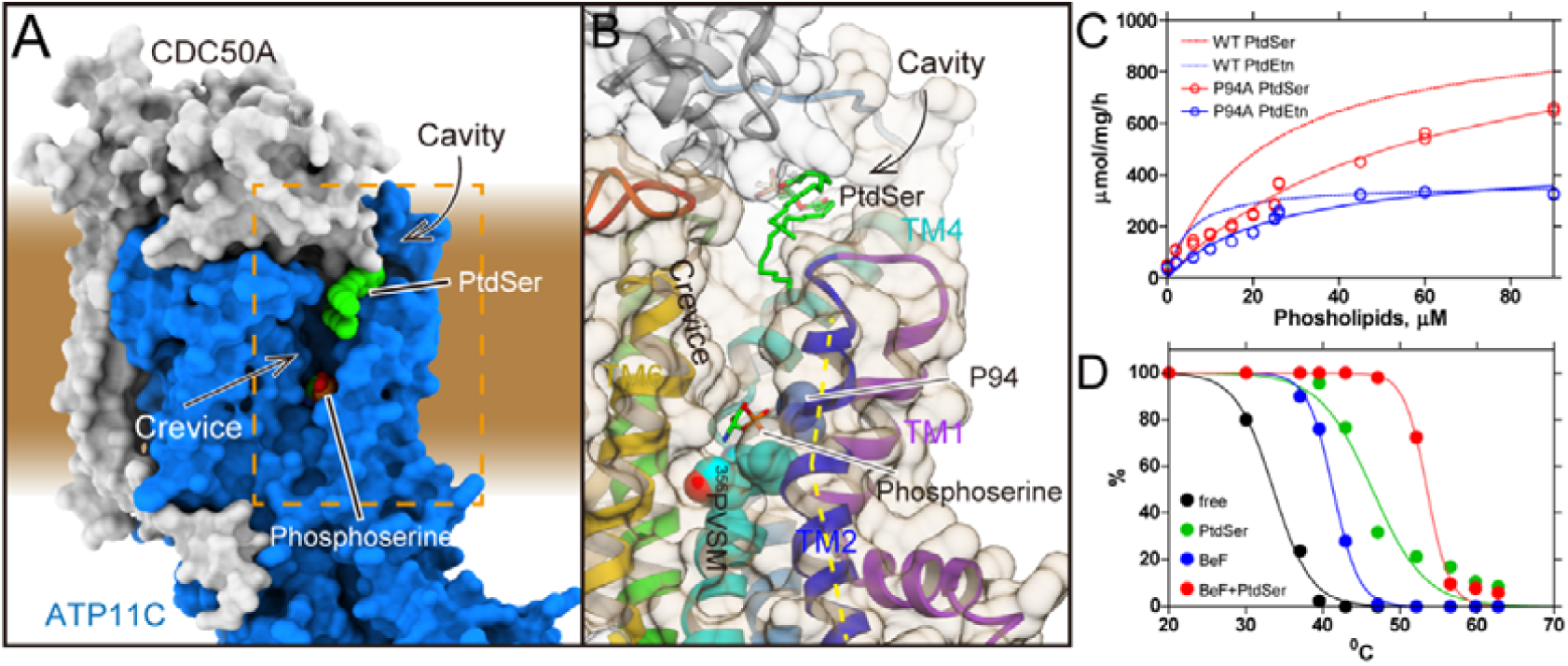
The crevice in the TM region. (A) Surface representation of the ATP11C-CDC50A complex shows the crevice in the TM region. Green spheres with CPK coloring represent phosphoserine at the occlusion site in the crevice, and PtdSer bound to the exoplasmic cavity. Surfaces of the atomic model of ATP11C and CDC50A are shown in blue and grey, respectively. (B) Close-up view of the membrane crevice indicated as a dotted box in A. The crevice is mostly composed of TM2, TM4 and TM6. Pro94 makes a kink at the exoplasmic side of TM2 (yellow dotted lines), which exposes the unwound region of TM4 (PVSM, shown in spheres) to the lipid bilayer phase. (C) PtdSer- or PtdEtn-dependent ATPase activities of wild-type (same as in Fig. 1) and Pro94Ala mutant as indicated in the figure. (D) Thermal stabilities of purified ATP11C-CDC50A complex determined by FSEC (see Methods). Peak values in the FSEC analysis were plotted as a function of treatment temperature in the absence (free) or presence of indicated substrates.

**Fig. 4.**
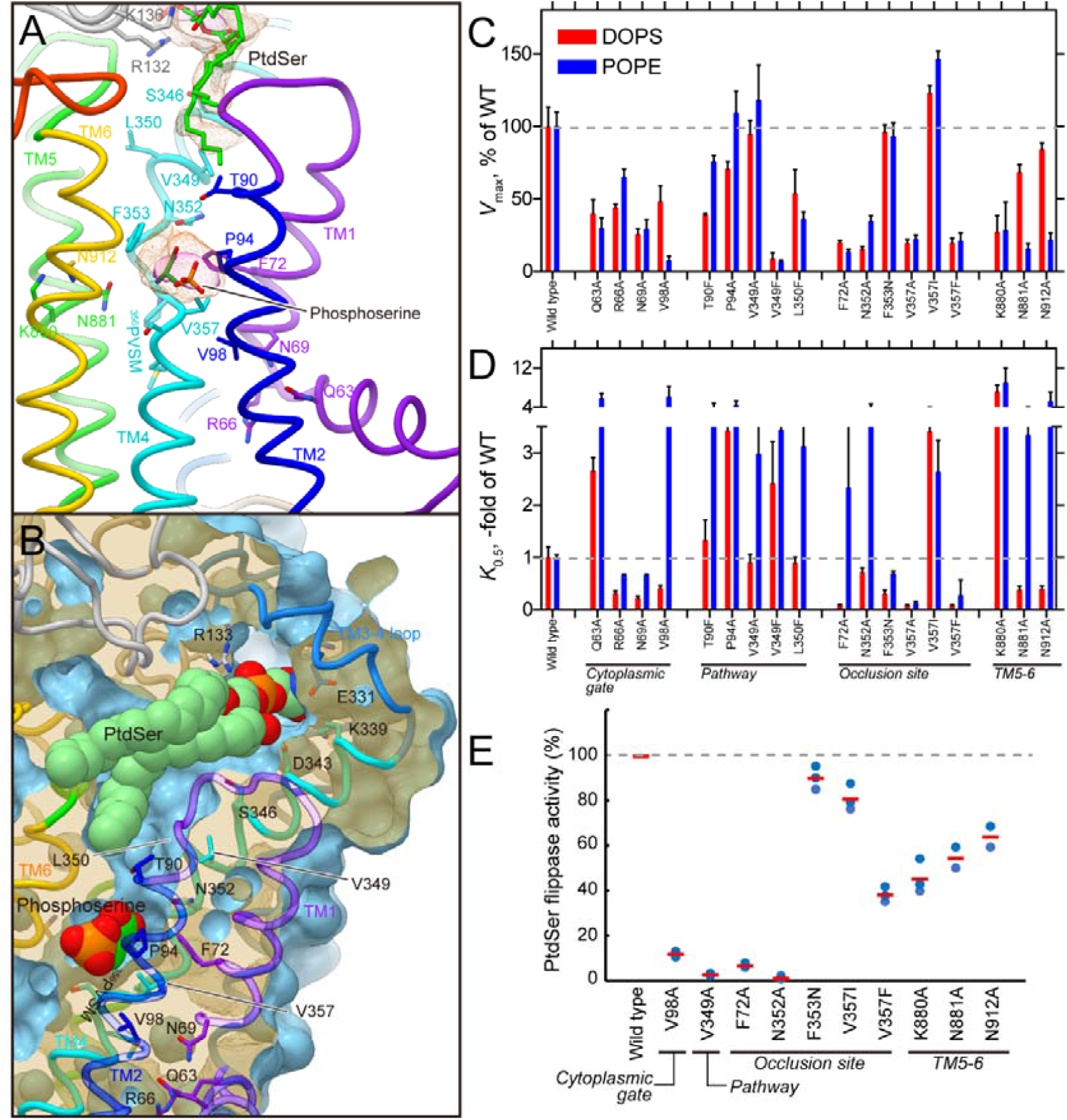
Transmembrane PtdSer occlusion site. (A) PtdSer occlusion site in detail, viewed from perpendicular to the membrane plane. Figure is displayed as in Fig. 3C. Phospho-L-serine (stick, green with CPK color) is modeled according to the observed electron density. (B) Phospholipid conduit along with TM4. Surface of the atomic model (light blue) is shown with superimposed ribbon model. Only surface model is clipped by the different plane at the position where TM4 is located, so as to show how the conduit runs along TM4. Its clipped surface is seen as transparent wheat color. Figure is drawn from TM1 and 2 viewpoint, with exoplasmic side-up. (C,D) *V*_max_ and *K*_0.5_ for indicated mutants determined from their PtdSer- or PtdEtn-dependent ATPase activities, as described in Fig. 3. (E) The PtdSer flippase activity. *ATP11A-ATP11C* knock-out (*DKO*) T-lymphoma cells expressing wild-type ATP11C or indicated mutants were incubated with 1 μM NBD-PS for 3 min. Experiments were performed two or three times, and flippase activity for NBD-PS is shown as a percentage to that of wild-type ATP11C. Horizontal red bars denote averages.

Well-ordered crystals were generated in the presence of both BeF_x_ and DOPS. The thermal stability^39^ of the purified ATP11C-CDC50A complex in the presence of both BeF_x_ and PtdSer (*T*_m_ = 53.5°C) is markedly higher than those in the absence (*T*_m_ = 33.6°C), or presence of either BeF_x_ (*T*_m_ = 41.3°C) or PtdSer alone (*T*_m_ = 46.4°C) (Fig. 2D), indicating that the BeF_x_-bound form binds PtdSer, and the enzyme is likely accumulats in a distinct PtdSer-bound, but not occluded, E2P form in the crystal. This conclusion is supported by the fact that an excess of PtdSer is required for preservation of the crystals when harvested for X-ray diffraction studies (Methods), indicating that PtdSer dissociates from the binding site at low to zero concentration. In fact, the electron densities found in the crystal structure (Fig. 1C, Fig. 3) led us to model two PtdSer molecules in the strucrture.

**Fig. 3.**
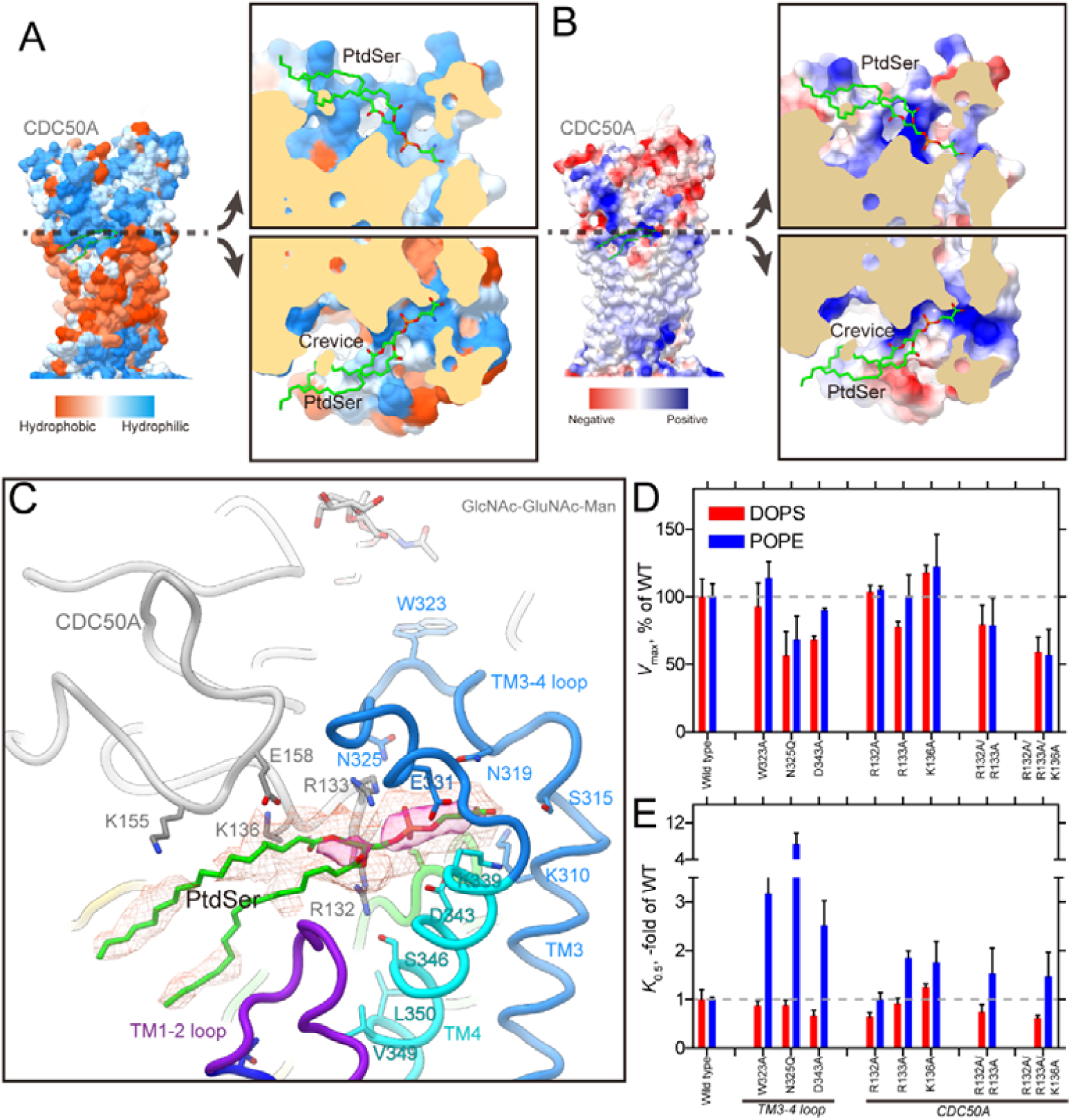
Exoplasmic cavity. (A,B) Surface representations of the atomic model of ATP11C-CDC50A according to their hydrophobicity (A) or Coulombic surface potentials (B). Each model was sliced at the plane where the phosphate moiety of PtdSer is located, along with the membrane plane, and opened up to show the surface of the exoplasmic cavity interior. Color codes as indicated in the figures. (C) PtdSer binding cavity in detail. Some hydrophilic amino acids surrounding PtdSer are shown as sticks. Color codes as in Fig. 1. Red mesh and transparent red surface represent the 2Fo-Fc electron density maps around the PtdSer molecule with contour levels of 1.0σ and 2.0σ, respectively. (D,E) The *V*_max_ and apparent affinity for PtdSer or PtdEtn derived from the ATPase activities of indicated mutants. The ATPase activity per mg of purified protein was determined and analyzed in the presence of varying concentrations of PtdSer or PtdEtn (Representatives shown in Fig. S7). The *V*_max_ is shown as a percentage of the wild-type value (D). The apparent affinities for phospholipids are expressed as the concentration giving half-maximum activation (*K*_0.5_), and plotted as x-fold of wild-type values (E).

### PtdSer binding at the exoplasmic cavity

There is a cavity at the exoplasmic side of the conduit (Fig. 3), which is formed by the TM3-4 loop of ATP11C and the exoplasmic domain of CDC50A around the surface of the membrane outer leaflet. This cavity is connected to the crevice observed in the TM region (Fig. 3, Fig. S6). Unexpectedly, the omit map showed an extra density at this position, and one PtdSer can be modeled here (Fig. 3C). The head group of the PtdSer is accommodated deep in the cavity, the surface of which is highly hydrophilic and electro-positive because of the many basic amino acids from both ATP11C and CDC50A arranged here (Fig. 3AB, Fig. S4). A cluster of basic amino acids is also observed in the cryo-EM structure of ATP8A1^31^. This positively-charged, and hydrophilic binding cavity would be favorable for attracting the negatively-charged phosphate group of a phospholipids. The hydrocarbon tails of the bound PtdSer lie along the membrane plane, projecting towards the membrane lipid phase. Single replacements of the positively-charged amino acids located at the entrance of the cavity produced only moderate effects on PtdSer- or PtdEtn-dependent ATPase activities, probably due to the number of remaining basic amino acids. This interpretation is supported by the approximately 50% reduction in the *V*_max_ of the triple mutant (Arg132Ala^CDC^/ Arg133Ala^CDC^/ Lys136Ala^CDC^) relative to that of wild-type (Fig. 3D). Beside these positively-charged amino acids, mutations in other hydrophilic amino acids in this cavity (Asn325Gln, Asp343Ala in ATP11C) produce a large reduction in apparent affinity for PtdEtn (Fig. 3E). Structural rigidity of the TM3-4 loop may also be important, because the apparent affinity for PtdEtn is also reduced in the alanine replacement of Trp323, which tethers the TM3-4 loop to the exoplasmic domain of CDC50A (Fig. 1D, Fig. 3C). These structural and functional data therefore suggest that this cavity is a priming site, responsible for the initial incorporation of phospholipids from the membrane outer leaflet – the beginning of the transport pathway.

The TM3-4 loop in ATP11C is 6 or 5 amino acid longer than that of ATP8A1 and Drs2p, respectively, and less conserved compared with other parts of the protein (Fig. S4). However, the hydrophilic and electro-positive nature of this cavity is essentially the same, at least, for the three flippases whose structures have been determined. Interestingly, some of mutations in the exoplasmic cavity had differing effects on the apparent affinities for PtdSer and PtdEtn (Fig. 3D,E), suggesting that this region does indeed associates with the transport ligands. This observation is consistent with a previously reported chimeric study on yeast flippase Dnf1, in which amino acids in the TM3-4 loop are shown to contribute to the ligand specificity^26^.

### PtdSer occlusion site

We also identified another PtdSer at the canonical occlusion site in the middle of the membrane (Fig 4), the position of which is close to that of PtdSer occluded in the ATP8A1 E2-P_i_ transition state. We modeled only a part of PtdSer (phospho-L-serine moiety) at this position according to the observed electron density (Fig. 4A), the other part including its acyl chains are likely disordered in the bulk lipid phase. Like other flippases as well as cation-transporting P-type ATPases, the conserved proline (Pro356) in the PVSM sequence (corresponding to a PISL motif in most of the other P4-ATPases) gives a characteristic unwinding at the middle of TM4, enabling the accommodation of a PtdSer head group at this position. The conduit ends at Val357 in the PVSM motif (Fig. 4B), which has been implicated as a gating residue for the phospholipids against the cytoplasmic inner leaflet^27^, similar to the glutamate residue in the corresponding PEGL motif of P2-type ATPases^40,41^. Hydrophobic residue of Val98 (corresponding to Ile115 in ATP8A2) supports the gating residue Val357 on its cytoplasmic side, and these hydrophobic amino acids form a tight seal as a cytoplasmic gate, which prevents the penetration of the PtdSer head group to the cytoplasmic side in the outward-facing E2P state. In fact, mutation of these amino acids severely impaired PtdSer- or PtdEtn-dependent ATPase activity as well as PtdSer transport activity in the plasma membrane (Fig. 4C-E), consistent with previous predictions for other flippases^26^. Mutation in the hydrophilic amino acids located at the cytoplasmic side of TM1 (Gln63Ala, Arg66Ala, Asn69Ala (corresponds to Asn220 in yeast flippase Dnf1)^42^) also showed reduced *V*_max_ values for ATPase activity relative to wild-type. These amino acids may contribute to the rigidity of the cytoplasmic gate, or are actually part of the conduit for the inner leaflet when the cytoplasmic gate opens. Val357 in ATP11C corresponds to Ile357 in ATP8A1, Ile364 in ATP8A2 and Ile508 in Drs2p, thus of almost all the P4-ATPases only ATP11A and ATP11C have a valine residue in this position (methionine in ATP8B3). In the case of ATP11C, mutation Val357Ile produces a remarkable reduction in apparent affinity for PtdSer and PtdEtn, while *V*_max_ is a bit higher than that of wild-type (Fig. 4C). In fact, the transport activity of Val357Ile is comparable to that of wild-type (Fig. 4E). In contrast, the ATPase activities of Val357Ala and Val357Phe, and the transport activity of Val357Phe are significantly reduced compared with those of wild-type. Evidently, correct size is important for gating residue Val357. Mutagenesis studies also reveal the important contribution of the conserved residues around the occlusion site (Phe72, Asn352) and TM5-6 (Lys880, Asn881 and Asn912), all alanine mutants showed significant reductions in either *V*_max_ or apparent affinity for PtdSer and/or PtdEtn determined through ATPase activity profiles, and transport activities as well, in good agreement with previous studies of ATP8A2^27,43^. Interestingly, Phe343 is not conserved among P4-ATPases, despite its close position to the phospholipid head group. In other PtdSer-transporting P4-ATPases, including ATP8A1, ATP8A2 and Drs2p, Phe343 in ATP11C is replaced with asparagine (Fig. S4), and the hydrophilic residue actually contributes to PtdSer coordination in the ATP8A1 E2-P_i_ structure^31^. The Phe354Asn mutant in ATP11C showed significantly higher affinity for PtdSer and PtdEtn relative to wild-type while keeping its *V*_max_ of ATPase activity and PtdSer transport activity comparable to wild-type level. Therefore, a hydrophilic or smaller side chain in this position is favorable for the accommodation of the aminophospholipid head group in the occlusion site. It can be speculated that ATP11C has Val357 and Phe354 instead of the most conserved isoleucine and asparagine of other PtdSer-dependent flippases, to fine-tune the PtdSer and PtdEtn transport activity relevant to a plasma membrane flippase. As an important note, mutations Val98Ala, Phe72Ala, Asn881Ala and Asn912Ala increases the apparent affinity for PtdSer, but lower that for PtdEtn, relative to the wild-type (Fig. 4D), therefore these mutants were able to discriminate PtdSer and PtdEtn.

### A transport model

Two PtdSer molecules bound to both ends of the conduit found in the crystal structure (Figs. 1,3 and 4) led us to put forward a transport model for the flippase, which is distinct from other models proposed so far^26,27,44^ (Fig. 5). Translocation of the phospholipid, either PtdSer or PtdEtn is initiated by its binding to the positively-charged and hydrophilic cavity composed of the CDC50A exoplasmic domain and the ATP11C TM3-4 loop (Fig. 3, Fig. S7). The electro-positive and hydrophilic nature of the cavity likely attracts the head group of the phospholipid from the outer leaflet layer of the membrane. The exoplasmic cavity connects to the longitudinal crevice along TM4 (Fig. 2, Fig. S6). Therefore, once the phospholipid head group is incorporated into the exoplasmic cavity, it may diffuse along the crevice while keeping its hydrocarbon chains projecting to the hydrophobic bulk lipid. Val349 in TM4 projects into the conduit (Fig. 4AB). Its replacement with bulky phenylalanine (Val349Phe) severely impairs PtdSer- and PtdEtn-dependent ATPase activity as well as PtdSer transport activity relative to wild-type, in contrast to the moderate effects of alanine substitution (Fig. 4C-E). In addition, phenylalanine mutations of residues close to Val349 (Thr90Phe in TM2 and Leu350Phe in TM4) also significantly lowers ATPase activity relative to the wild-type. These amino acid residues are located between, and distant from, the two substrate binding sites and clearly bulky substitutions impede diffusion of the phospholipid head group from the exoplasmic cavity to the occlusion site. In the occlusion site, the phospholipid head group must be coordinated by conserved hydrophilic amino acids such as Asn352 and Asn881. The gating residue Val357 blocks further penetration of the phospholipid to the cytoplasmic inner leaflet. PtdSer binding to the occlusion site may be the signal to induce the conformational change required for reaching the PtdSer-occluded E2-P_i_ transition state. Some of the hydrophilic amino acids in the TM1-2 loop may join in the coordination of the head group, as observed in the ATP8A1 E2-P_i_ structure (Fig. S6)^31^. Movement of TM1-2, as a result of the loop involvement, adjusts the A domain to induce dephosphorylation of E2P. Attainment of the E2 or E1state, the next step, opens the cytoplasmic gate. A physical pathway from the occlusion site to the inner leaflet need to be opened. Mutation of amino acid residues around the cytoplasmic gate largely inhibited phospholipid dependent ATPase activity (Fig. 4). Movement of the TM1-2 helix bundle outwards may be needed for the phospholipid to move to the cytoplasmic inner leaflet, although this has not yet been elucidated.

**Fig. 5.**
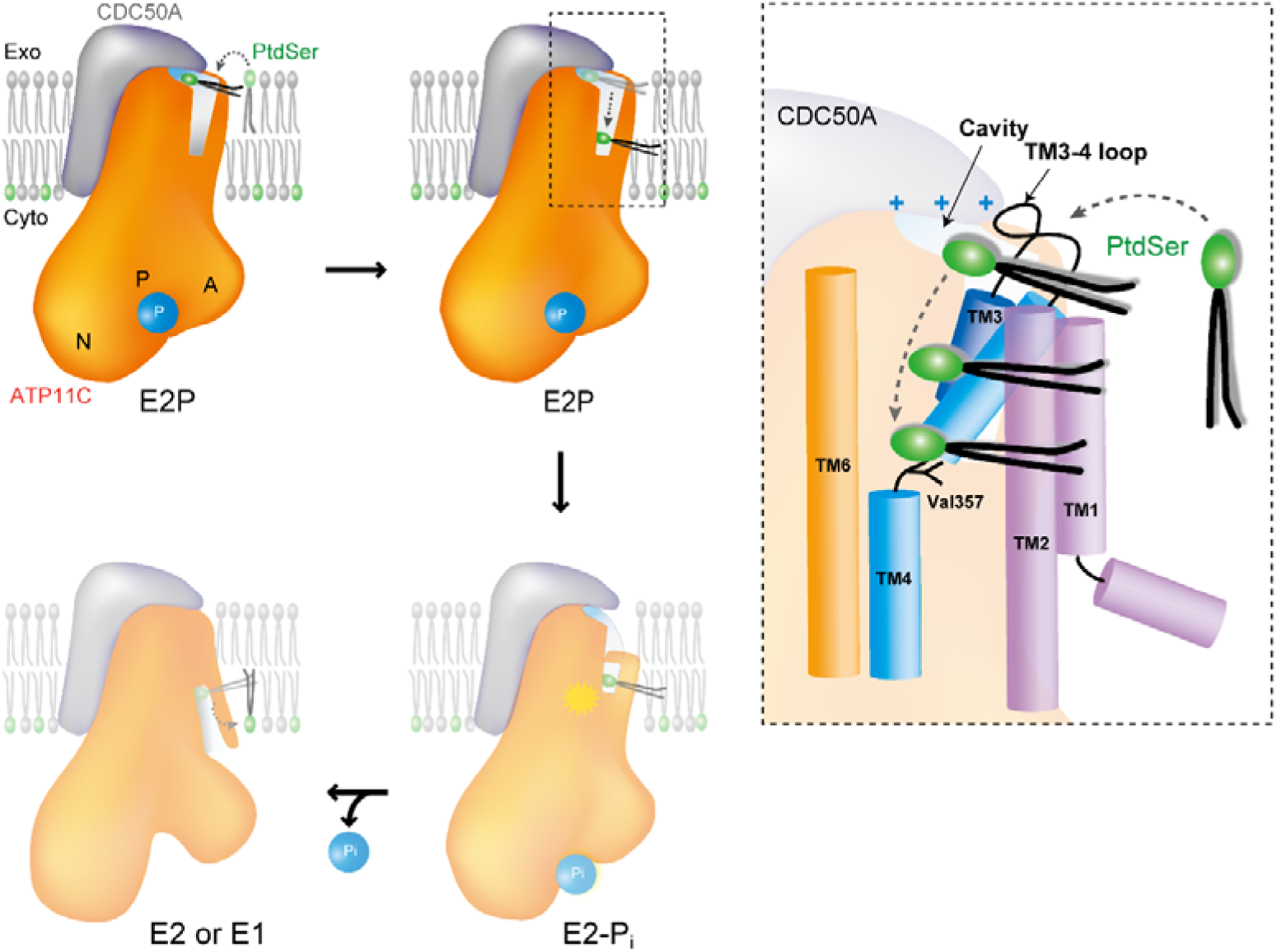
A phospholipid transport model for ATP11C. (A) Schematic drawing of the transport mechanism by ATP11C-CDC50A complex. In the outward-open E2P state (present structure), phospholipid enters from the surface of outer leaflet to the exoplasmic cavity by changing its orientation (upper left). Trapped phospholipid head group diffuses along the membrane crevice with its hydrophobic tails extending out to the hydrophobic core of the bilayer (upper right). When the phospholipid head group reaches Val357 in the middle of the membrane, the phosphate head group is occluded by closing the crevice with the TM1-2 helix bundle (lower right). This process is coupled with dephosphorylation of E2P as seen in the ATP8A1 E2-P_i_ transition state structure. Complete dephosphorylation may further open the cytoplasmic gate, and phospholipid is translocated to the cytoplasmic inner leaflet (lower left). (B) Close-up view of the membrane crevice indicated as dotted box in A. Phospholipid headgroup traverses from positively-charged exoplasmic cavity to the occlusion site near Val357 along with TM4.

Translocation of the phospholipid across the two leaflets is energetically extremely unfavorable due to the amphiphilic nature of phospholipids^45^, and the rate of spontaneous phospholipid flip-flop is order of several hour to several days. In our envisaged translocation mechanism, the most energy-consuming step may be moving the hydrophilic head group from the outer membrane surface to the transport conduit. In this step the phospholipid head group needs to disconnects from the polar interactions formed with neighboring phospholipids, surrounding membrane proteins as well as water molecules, and also needs to change its orientation approximately 90° from a vertical to horizontal orientation in the lipid bilayer. The structure and resulting transport model answer the fundamental question in the translocation mechanism of flippase -how phospholipid reaches the transport conduit from the outer leaflet. The hydrophilic surface of the exoplasmic cavity interior provides an environment similar to the water-facing membrane surface and lowers the energetic barrier required for acquisition of the phospholipid head group. Many membrane transport proteins seem to employ a common strategy for the translocation of their specific substrates; the environment of the substrate binding site of the protein mimics that of the location from whence the substrate comes. Flippases are not an exception, and this strategy is applied in the most energy-consuming step in the sequence of the lipid flipping process by the ATP11C-CDC50A complex.

## Acknowledgement

We thank M. Taniguchi for technical assistance, Dr. T. Nishizawa for sharing unpublished results of the ATP8A1 structure and Dr. D. McIntosh for improving the manuscript. This work was supported by Grants-in-Aid for the Scientific Research (17H03653), Basis for Supporting Innovative Drug Discovery and Life Science Research (BINDS) from Japan Agency for Medical Research and Development (AMED), Takeda Science Foundation (to K.A.); Core Research for Evolutional Science and Technology from JST (JPMJCR14M4, to S.N. and K.A.); Grants-in-Aid for Scientific Research (S), the Japan New Energy and Industrial Technology Development Organization (NEDO), and the Japan Agency for Medical Research and Development (AMED) (to Y.F.). This work is a part of a project (2018B2703 and 2019B2707) at SPring-8. This research is partly supported by the Platform Project for Supporting Drug Discovery and Life Science Research (BINDS) from AMED under Grant number JP18am0101070.

## Author Contributions

Y.F., S.N. and K.A. designed the study. H.N. and K.A. were responsible for protein expression. H.N. purified and crystallized the protein. H.N. K.S. and K.A. performed biochemical analysis. H.N., K.H. and K.A. collected X-ray diffraction data. K.H. merged X-ray diffraction data. K.I. and K.A. analyzed the structure. K.I. and K.A. interpreted the structure. H.N. K.S. and K.A. wrote the manuscript with agreement of all authors.

## Author Information

Atomic coordination and structure factors for the structures reported in this work have been deposited in the Protein Data Bank under accession number NXXX.

## Materials and Methods

### Protein expression and purification

Human *ATP11C* (NCBI: XM_005262405.1)^4^ was sub-cloned into a hand-made vector as described previously^13^. Both of the amino terminal 7 amino acids (ΔN7) and the carboxyl terminal 38 amino acids (ΔC38) of hATP11C were truncated, and the Flag epitope tag (DYKDDDDK), hexa-histidine tag, the enhanced green fluorescence protein (EGFP) followed by a tobacco etch virus (TEV) protease recognition sequence were attached to the amino terminal of the deletion mutant (ATP11C_cryst). Human *CDC50A* cDNA (NCBI: NM_018247.3) was sub-cloned into the vector independently. The Asn190Gln and Ser292Trp double mutation was introduced into the construct to regulate its glycosylation status (CDC50A_QW). The former mutation is simply aimed to prevent *N*-linked glycosylation. The latter is to eliminate *O*-linked glycosylation at Ser292 and at the same time to increase the efficiency of *N*-linked glycosylation at Asn294^48^, because these residues appeared to be glycosylated alternatively. The heterodimer composed of ATP11C_cryst and CDC50A_QW was successfully expressed in the plasma membrane using baculovirus-mediated transduction of mammalian Expi293 cells (Thermo) for 48 h at 31.5 °C as described previously^28,46^. The harvested cells were directly solubilized with 1.5 % (w/v) n-decyl β-D-maltoside in a lysis buffer containing 40 mM MES/Tris (pH 6.5), 200 mM NaCl, 2 mM Mg(CH_3_COO)_2_, 1 mM ATP, 1 mM dithiothreitol, 0.1 mM ethylene glycol-bis(2-aminoethylether)-*N,N,N’,N’*-tetraacetic acid (EGTA) and protease inhibitor cocktail (Roche) on ice for 20 min. After removing the insoluble material by ultracentrifugation (200,000×g for 1h), the supernatant was mixed with anti-Flag M2 affinity resin (Sigma-Aldrich) for 1 h at 4 °C. The resin was washed with 20 column volumes of buffer consisting of 20 mM MES/Tris (pH 6.5), 200 mM NaCl, 5% (v/v) glycerol, 1 mM Mg(CH_3_COO)_2_, 0.1 mM ATP, 0.1 mM EGTA and 0.03% octaethylene glycol monododecyl ether (C_12_E_8_, Nikko Chemical). Flag-EGFP tagged ATP11C was eluted with 0.2 mg/ml Flag peptide (Sigma-Aldrich) in the wash buffer. Eluted proteins were incubated with TEV protease and MBP-fusion endoglycosidase (EndoHf, New England Biolabs) at 4 °C overnight. Released affinity tags containing Flag-EGFP were removed from the mixture by a Ni-NTA resin (QIAGEN). The non-absorbed fractions were concentrated and subjected to a size-exclusion column chromatograph using a Superose6 Increase column (GE Healthcare), equilibrated in a buffer comprising 20 mM MES/Tris (pH 6.5), 1%(v/v) glycerol, 50 mM NaCl, 5 mM MgCl_2_, and 0.03% C_12_E_8_. Peak fractions were collected and concentrated to 10 mg/ml. See Fig. S1 for the purity of the sample at each step. The concentrated ATP11C samples were mixed with 1 mM ADP, 0.5 mM BeSO_4_, 1.5 mM NaF, and 0.1 mg/ml dioleoyl phosphatidylserine (DOPS), and added to the glass tubes in which a layer of dried dioleoyl phosphatidylcholine (DOPC) had formed, in a lipid-to-protein ratio of 0.2. C_12_E_8_ was added to the glass tubes in a protein-to-detergent ratio of 0.5 to 2.0, and the mixture was incubated overnight at 4 °C in a shaker mixer operated at 120 rpm^29^. After removing the insoluble material by ultracentrifugation, lipidated samples were subjected to crystallisation. Note that the effect of truncation of the both terminal amino acids of ATP11C, the double mutation introduced to CDC50A and deglycosylation of CDC50A on the PtdSer- and PtdEtn-dependent ATPase activity were negligible compared with wild-type without endoglycosidase treatment.

### Gene editing for CDC50A

The CRISPR (Clustered Regularly Interspaced Short Palindromic Repeats)-Cas (CRISPR-associated) system with pX330 vector (Addgene) was used to edit the CDC50A gene in HEK293S GnT1-cells as described ^4^. The sgRNA sequences for human CDC50A were as follows; 5’-CACCG<u>GGCAACGTGTTTATGTATTA</u>-3’ and 5’-AAAC<u>TAATACATAAACACGTTGCC</u>C-3’. Target protospacer sequences are underlined.

### Crystallization and data collection

Crystals were obtained by vapour diffusion at 20°C. The lipidated 10 mg/ml protein sample obtained either from Expi293 cell or CDC-KO cells, containing 1 mM ADP, 0.5 mM BeSO_4_, 1.5 mM NaF, and 0.1 mg/ml DOPS was mixed with reservoir solution containing 10% (v/v) glycerol, 14-17% PEG4000, 0.4 M MgSO_4_, and 2 to 5 Mm β-mercaptoethanol. Crystals made using protein samples purified from Expi293 cells grew in a thin plate-like shape with the dimensions of 800 × 500 × 50 μm in 2 weeks. In contrast, crystals from CDC-KO cells grew as small crystals usually less than 50 μm with a polyhedron shape. These crystals were picked up with LithoLoops (Protein Wave Corporation), and flash frozen in liquid nitrogen. The crystals were harvested in the presence of 10% (v/v) glycerol, 14-17% PEG4000, 0.4 M MgSO_4_, 4 mg/ml DOPS, 20 mM MES/Tris (pH 6.5), 50 mM NaCl, 5 mM MgCl_2_, 5 mM β-mercaptoethanol, 2% C_12_E_8_, 1 mM ADP, 0.5 mM BeSO_4_ and 1.5 mM NaF. Note, when crystals were harvested in the absence of DOPS, a few percent approximately of crystals gave X-ray diffraction better than 4Å. In the presence of 4 mg/ml DOPS, however, the number of well-diffracting crystals mostly increased to approximately 30% of all harvested crystals. Despite the different crystal morphologies of the crystals obtained from Expi293 cells and CDC50A KO cells (Fig. S1), the crystals show the same space group (*P*2_1_2_1_2_1_) and unit cell size (*a* = 100.5Å, *b* = 232.8Å, *c* = 492.9 Å, α = β = γ = 90°).

X-ray diffraction data were collected at the SPring-8 beamline BL32XU, BL41XU and BL45XU. For the large plate-like crystals obtained from Expi293 cells, X-ray diffraction data were collected by helical scan method^49^, or by irradiating micro-focus beam from the direction perpendicular to the *c*-axis by monitoring crystals on 90° bent LithoLoop (Fig. S1). Crystals from CDC-KO cells were too small to collect full data set from each crystal, and well-diffracted crystal could not be determined from its morphology. Therefore, multiple crystals were mounted on a *ϕ*1 μm LithoLoop, and the raster scan was performed to identify well-diffracted crystals. After selecting target crystals, 10° small-wedge data were collected from each individual crystal (Fig. S1). Total 1,588 crystals were used for the data collection and some of them were performed automatic manner by using ZOO system^50^.

### Structural determination and analysis

All the diffraction data from individual 1,588 well-diffracting crystals were processed and merged using automatic data processing system KAMO^47^ with XDS^51^. Structure factors were subjected to anisotropy correction using the UCLA MBI Diffraction Anisotropy server^52^ (http://services.mbi.ucla.edu/anisoscale/). The structure was determined by molecular replacement with PHASER, using a homology model based on the cryo-EM structure of the E2P state of ATP8A1 (PDB ID: 6K7L) as a search model. Coot^53^ was used for cycles of iterative model building and Refmac5^54^ and Phenix^55^ were used for refinement. The final crystallographic model of BeF_x_-bound human ATP11C at a 3.8Å resolution, refined to *R*_work_ and *R*_free_ of 0.29 and 0.36 was deposited in the PDB with accession code PDB: 6LKN. Figures were prepared using UCSF Chimera^56^ and PYMOL (https://pymol.org).

### Activity assay using recombinant proteins

To measure the ATPase activity, Flag-EGFP tag connected by the TEV cleavage site to the N-terminal tail of ATP11C_cryst was used to estimate its expression level by fluorescence size-exclusion column chromatography (FSEC)^57^. The original and additional mutant complexes of ATP11C_cryst and the CDC50A_QW were expressed using the BacMam system and purified in a smaller batch format as described above except for TEV protease digestion and endoglycosidase treatment. The purified proteins (the purity of samples used for the ATPase measurement was comparable to lane 4 of SDS-PAGE in Fig. S1) were subjected to an ATPase activity assay as described previously^58^. Briefly, partially purified ATP11C (wild-type or mutants) was suspended in buffer comprising 40 mM HEPES, 2 mM MgCl_2_, 2 mM ATP, 2% glycerol, 100 mM NaCl, 0.03 mg/ml C_12_E_8_ (pH 7.0 adjusted by Tris) and indicated concentrations of phospholipids (DOPS or POPE, dissolved as 10 mg/ml stock in 2% C_12_E_8_), in 96-well plates. Reactions were initiated by incubating the samples at 37 °C using a thermal cycler, and maintained for 1h. Reactions were terminated, and the amount of released inorganic phosphate was determined colorimetrically using a microplate reader (TECAN). Samples used for the ATPase measurement were analyzed by FSEC with monitoring with Trp fluorescence (Ex 280 nm, Em 320 nm) monitoring, and peak fluorescence values were determined. The peak values of samples were compared to that of a fully purified sample used for the crystallization whose protein concentration was accurately determined by UV absorption, and protein concentrations for each measured sample were estimated. The phospholipid concentration-dependent ATPase activities were plotted, and data fitted as described previously^43^ to estimate apparent affinities (*K*_0.5_) and *V*_max_ using PRISM4 software. For all measurements, data were duplicated at twelve different phospholipid concentrations for a single measurement, and at least three independent measurements were conducted for each mutant. The *K*_0.5_ and *V*_max_ values plotted in Fig. 3 and 4 are mean and S.D.s estimated from at least three independent measurements, and representative ones are shown in Fig. S5. Note, the PtdSer- or PtdEtn-dependent specific ATPase activities of ATP11C_cryst-CDC50A_QW complex were almost the same as those of wild-type. We therefore refer to ATP11C_cryst-CDC50A_QW as wild-type in the activity assay for simplicity.

### Flippase assay

Flippase activity was determined as described^59^. In brief, *ATP11A-ATP11C* double-deficient WR19L mutant (*DKO*) cells were transformed with retroviruses carrying cDNA for human FLAG-tagged CDC50A and EGFP-tagged wild-type ATP11C or mutants. The stable transformants were then subjected to cell sorting for EGFP with FACSAriaII (BD Biosciences), cells at the same EGFP-intensity were sorted, and expanded. Amounts of EGFP-tagged proteins and their localization to the plasma membrane were verified by Western blotting and a confocal microscope (FV-1000D; Olympus), respectively. For the flippase assay, *DKO* cells and its transformants expressing wild-type ATP11C or mutants were incubated with 1 μM NBD-PS (1-oleoyl-2-{6-[(7-nitro-2-1,3-benzoxadiazol-4-yl)amino]hexanoyl}-*sn*-gylcero-3-phosp hoserine for 3 min at 20 °C in Hanks’ balanced salt solution (HBSS) containing 1 mM MgCl_2_ and 2 mM CaCl_2_. The cells were collected by centrifugation, resuspended in HBSS containing 5 mg/ml fatty acid-free BSA to extract nonincorporated lipids, and analyzed by FACSCanto II (BD Biosciences).

### Thermal stability

Purified samples were incubated at the indicated temperatures for 10 min in the presence of 40 mM HEPES, 100 mM NaCl, 2 mM MgCl_2_ (free) with 1 mM BeSO_4_, 3 mM NaF (BeF) and/or 0.1 mg/ml DOPS (PtdSer). After incubation, samples were cooled on ice and denatured aggregates removed using a membrane filter (pore size 0.22 μm), and the resulting filtrates were analyzed by size-exclusion column chromatography using Superose 6 Increase 10/150 GL (GE healthcare). Peak values at the retention time for the complex were plotted as a function of incubation temperature, and their *T*_m_ values estimated.

---

**Fig. S1.**
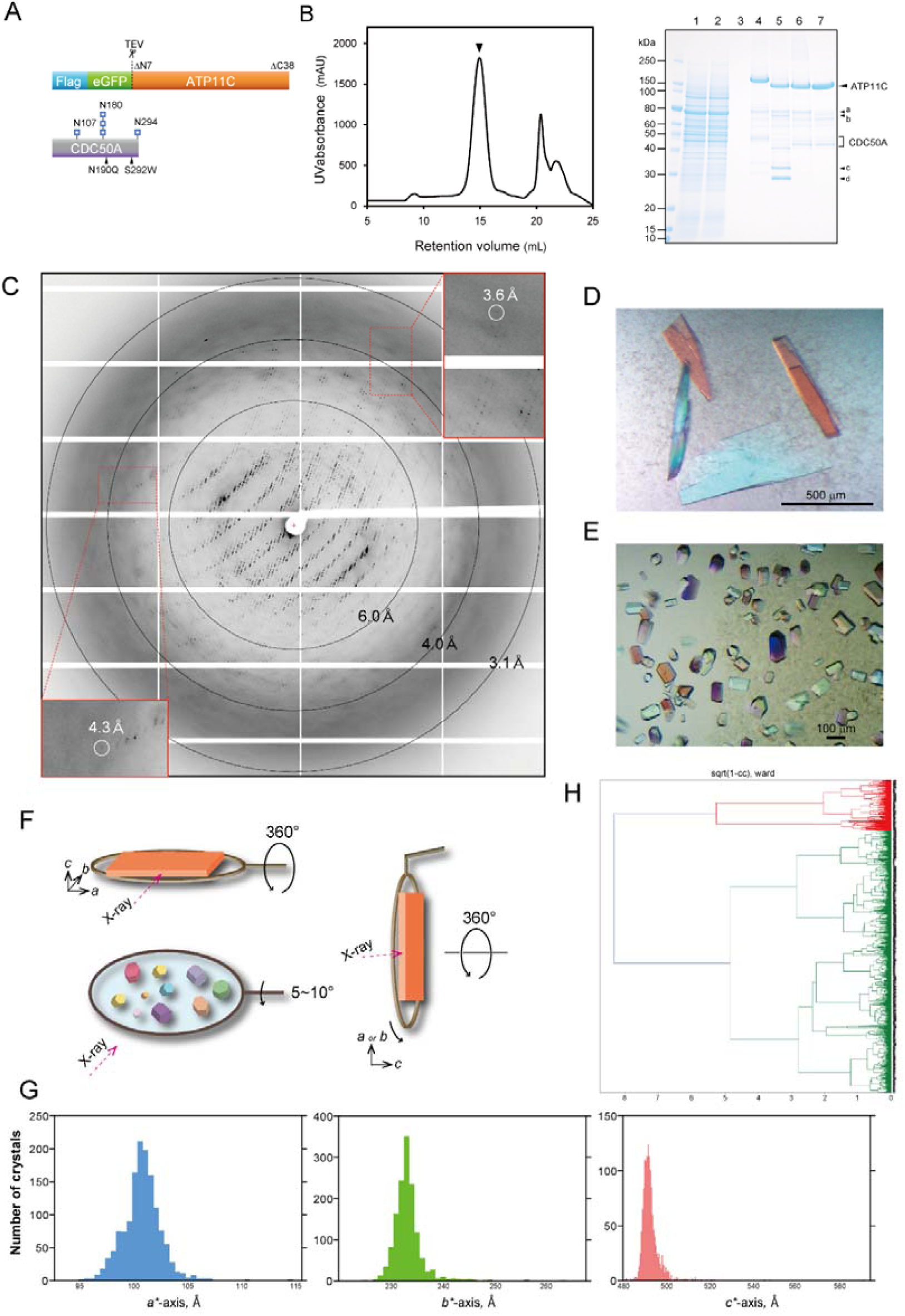
Structural determination of ATP11C-CDC50A complex. (A) Construction of TP11C and CDC50A used in crystallization. See Methods for details. (B) Purification of ATP11C-CDC50A complex. Lane 1: solubilized cell lysate, lane 2: pass-through of Flag resin, lane 3: wash fraction, lane 4: elution by Flag peptide (subjected to ATPase assay), lane 5: TEV proteinase- and endoglycosidase-treated sample, lane 6: pass-through fraction of Ni-NTA and amylose resin, lane 7: concentrated peak fractions by size-exclusion chromatography (arrowhead in the left panel). Arrowheads on the right indicated as follows, a: HSP70, b: EndoHf, c: cleaved eGFP, d: TEV proteinase. The elution profile of ATP11C-CDC50A complex by size-exclusion column chromatography is shown on the left. (C) Representative X-ray diffraction obtained from a plate-like crystal shown in D. Diffraction spots better than 3.6Å were obtained along the *c*^*^-axis, whereas these are limited to around 4∼6Å in directions along with *a*^*^- and *b*^*^-axes, thus strongly anisotropic. (D,E) Three-dimensional crystals obtained from the samples purified from Expi293 cells, showing thin, but large plate-like crystals (D). In contrast, small crystals were obtained from CDC50A-KO cells (E). (F) Data acquisition strategy. We employed normal type LithoLoops for helical scan data acquisition from large single crystals. However, because crystals showed strong anisotropy, we also collected data sets by irradiating X-ray beam from the direction perpendicular to the *c*-axis by using 90° bent type LithoLoops. For the small but well diffracting crystals obtained from CDC50A-KO cells, data from each individual crystals was collected for 10°. All of these crystals showed identical unit cell size and symmetry regardless of crystal morphologies and expression cell types, as seen in the histograms of unit cell dimensions (G). All diffraction data from 1,588 crystals were finally merged into a single data set (H), and used for the molecular replacement.

**Fig. S2.**
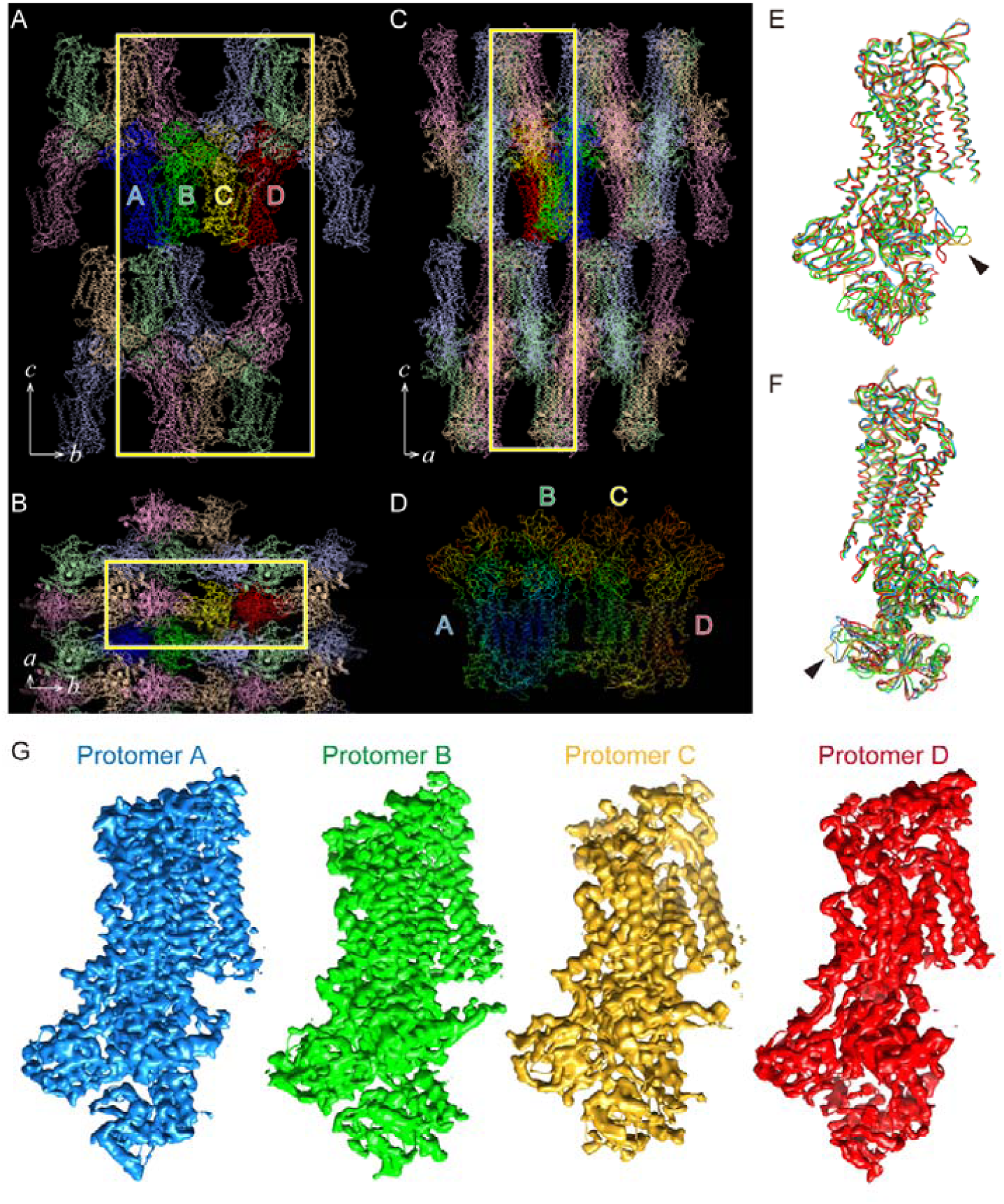
Crystal packing. (A-C) An asymmetric unit contains four protomers (A, B, C and D) shown in blue, green, yellow and red, respectively. Their symmetry-related molecules are shown in light colors. Yellow boxes indicate unit cells view from different direction in A-C as indicated in the figures. (D) Molecules in the asymmetric unit are displayed according to their temperature factors. Colors gradually change from blue (16) to red (283). (E,F) Comparison of the molecular conformation of four protomers. Arrowheads indicate loop structures, the conformations of which are variable among the four protomers. (G) 2F_o_-F_c_ electron density maps of the four protomers at the same contour level of 1.5σ.

**Fig. S3.**
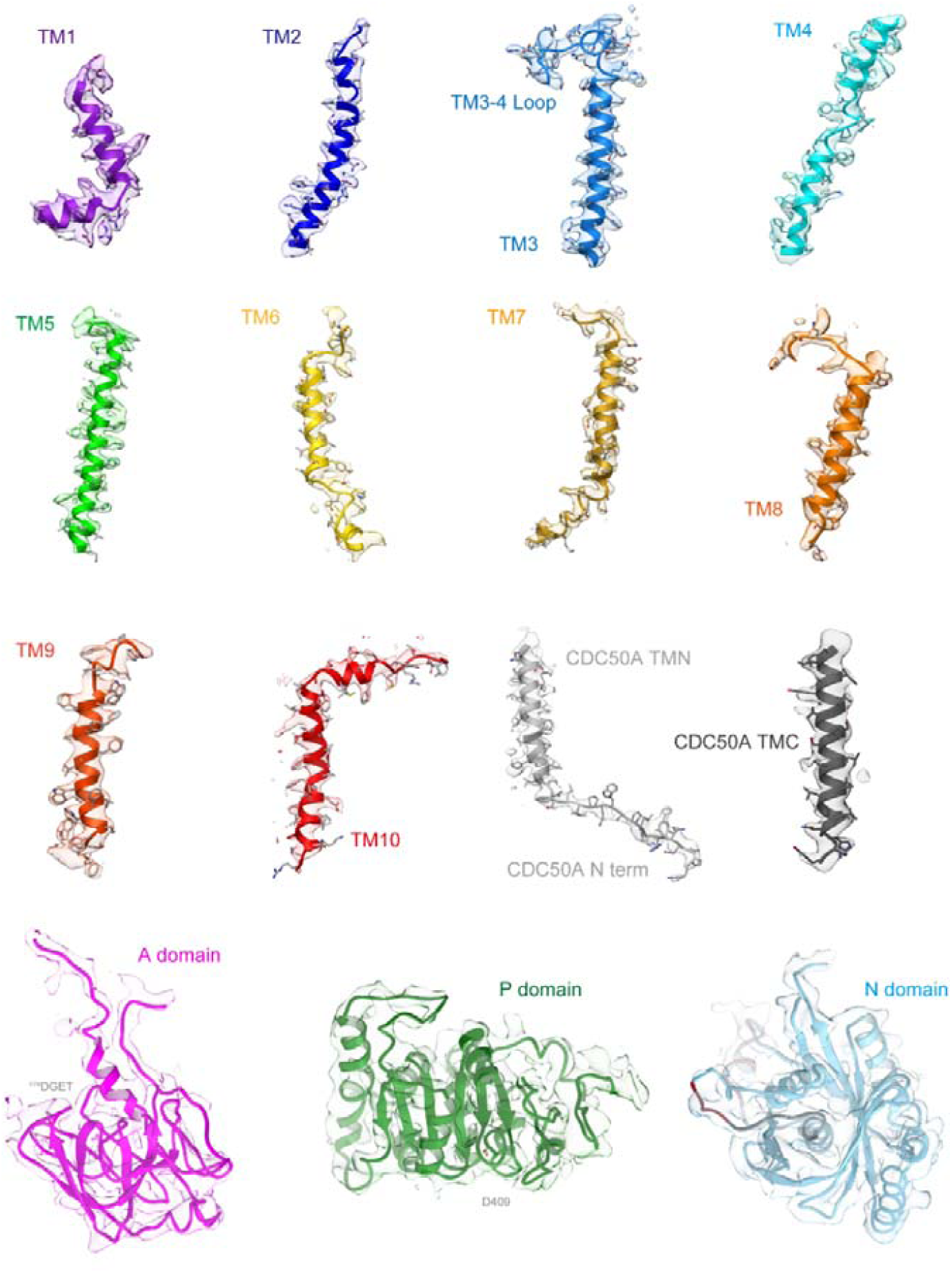
Electron density maps. Surface represents 2F_o_-F_c_ electron density maps of the indicated regions with 1.5σ contour level. Color code as in Fig. 1.

**Fig. S4.**
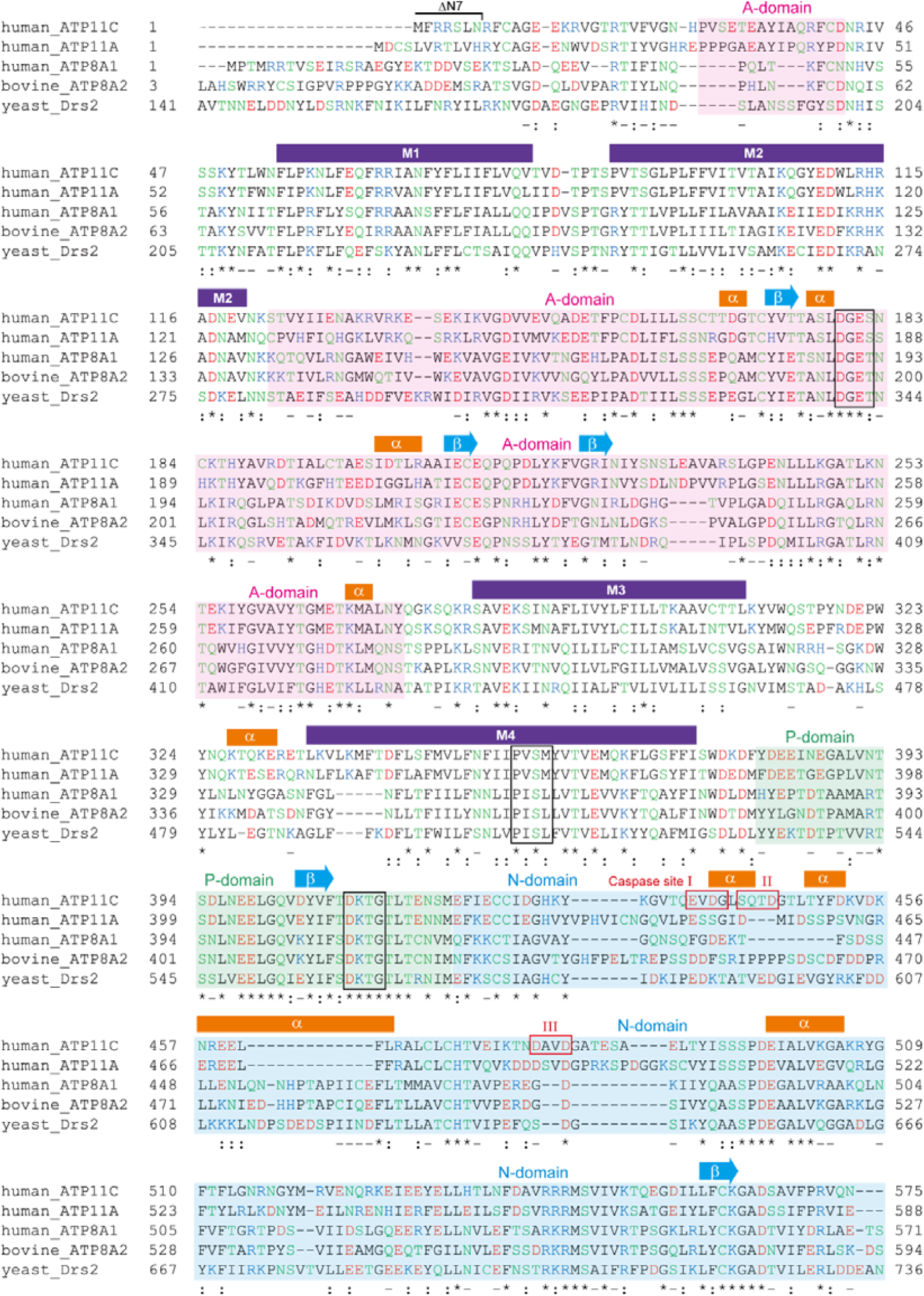
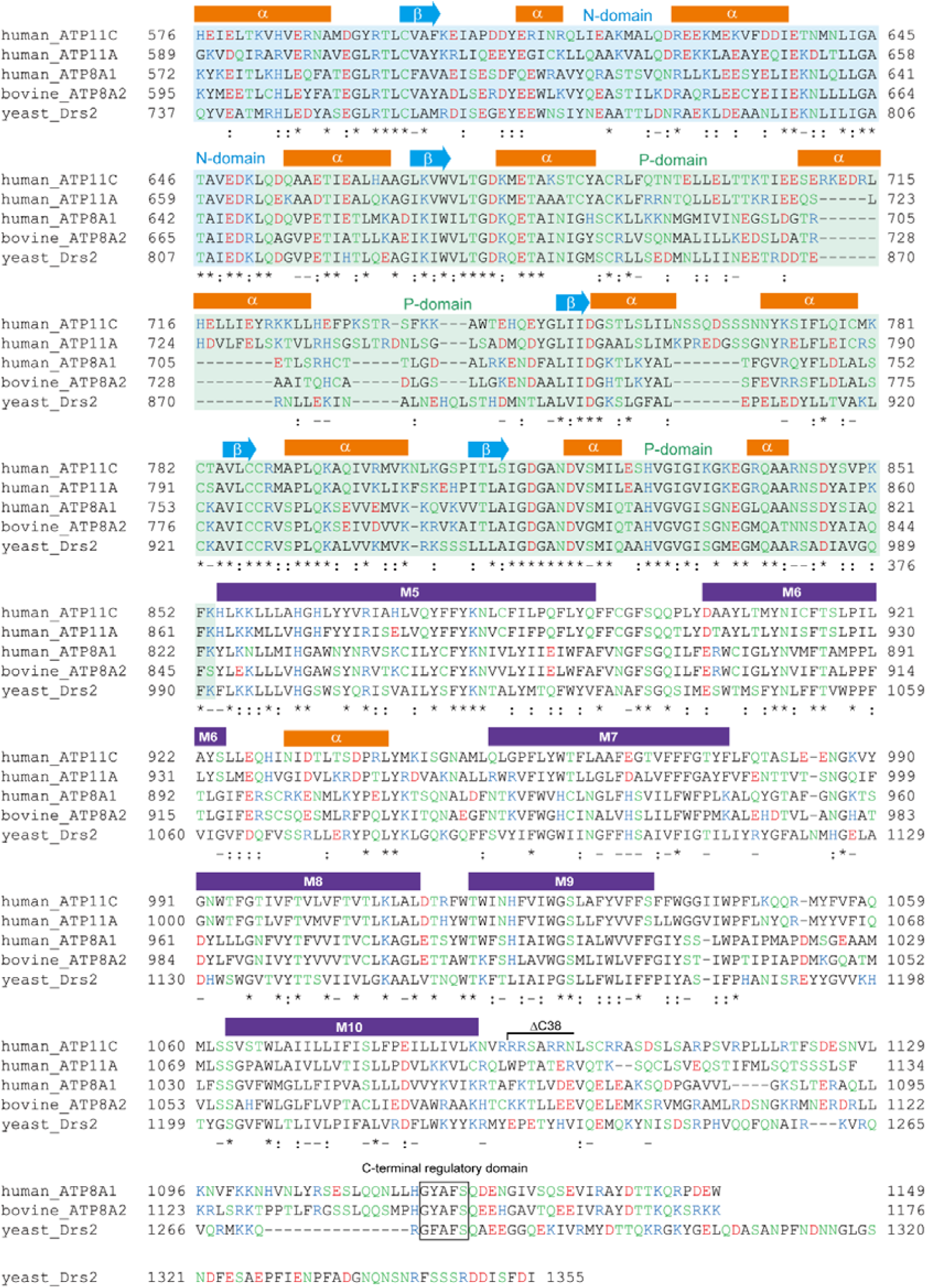
Sequence alignment. Primary sequences of indicated P4-ATPases isoforms were aligned using software MAFFT ver.7^35^ and large gaps introduced in the non-conserved region were manually edited. Cytoplasmic domains (A, pink; P, green; N, light blue), secondary structures (α-helices, β-sheets and TM helices) and mutations introduced for the crystallized construct (ΔN7, ΔC38) are indicated above the alignment. The degree of conservation among evaluated sequences is indicated below the alignment. Acidic, basic, hydrophilic or hydrophobic amino acids are indicated as red, blue, green and black characters, respectively. Gene and protein ID is as follows; human ATP11C (NCBI:_XM_005262405.1), human ATP11A (Uniprot: P98196-1), human ATP8A1 (UniProt: Q9Y2Q0-2), bovine ATP8A2 (Genebank: GQ303567.3), and Saccharomyces cerevisiae Drs2 (UniProt: P39524-1).

**Fig. S5.**
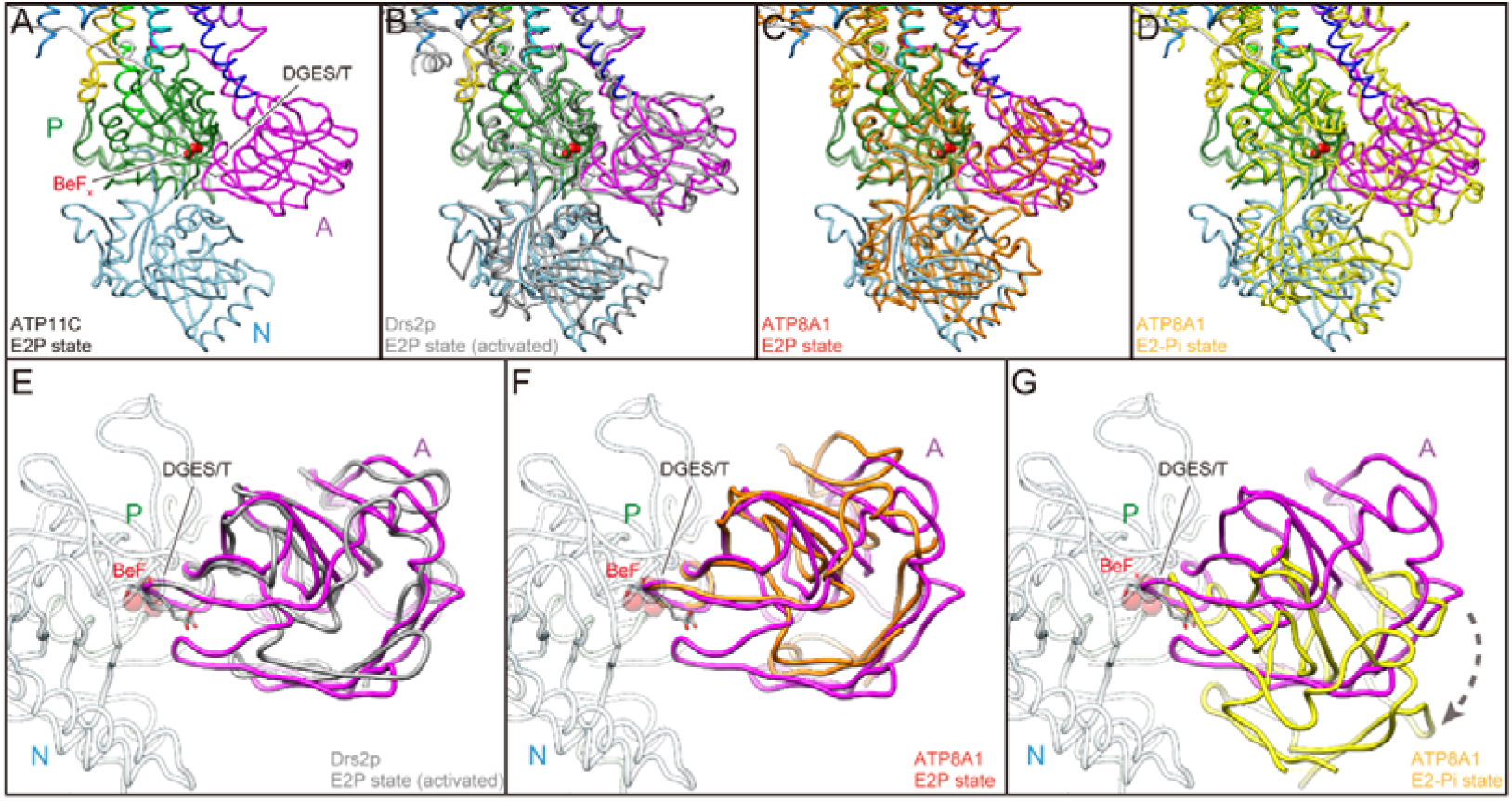
Cytoplasmic domains. (A-D) Comparison of the relative orientation of the cytoplasmic domains viewed along with the membrane plane. Cytoplasmic domains of ATP11C are shown in worm models with the same colour code as in Fig. 1C (A). Atomic models of Drs2p in E2P activated form (B, grey), ATP8A1 E2P form (C, orange) and ATP8A1 E2-P_i_ form (D, yellow) are superimposed on the ATP11C structure according to their P domain structures to show relative orientations of A and N domains. (E-G) Azimuthal position of the A domain is compared. A domain of Drs2p (E), ATP8A1 E2P state (F) and E2-P_i_ state (G), and these models are superimposed on the ATP11C structure (only A domain is highlighted in magenta, and others are shown in transparent colors) as in A-D. Dotted arrow indicates the different azimuthal positions between ATP11C E2P state and ATP8A1 E2-P_i_ state. Phosphate analogue BeF_x_ (red spheres) and DGES/T motif (sticks) in ATP11C structure are highlighted in all figures.

**Fig. S6.**
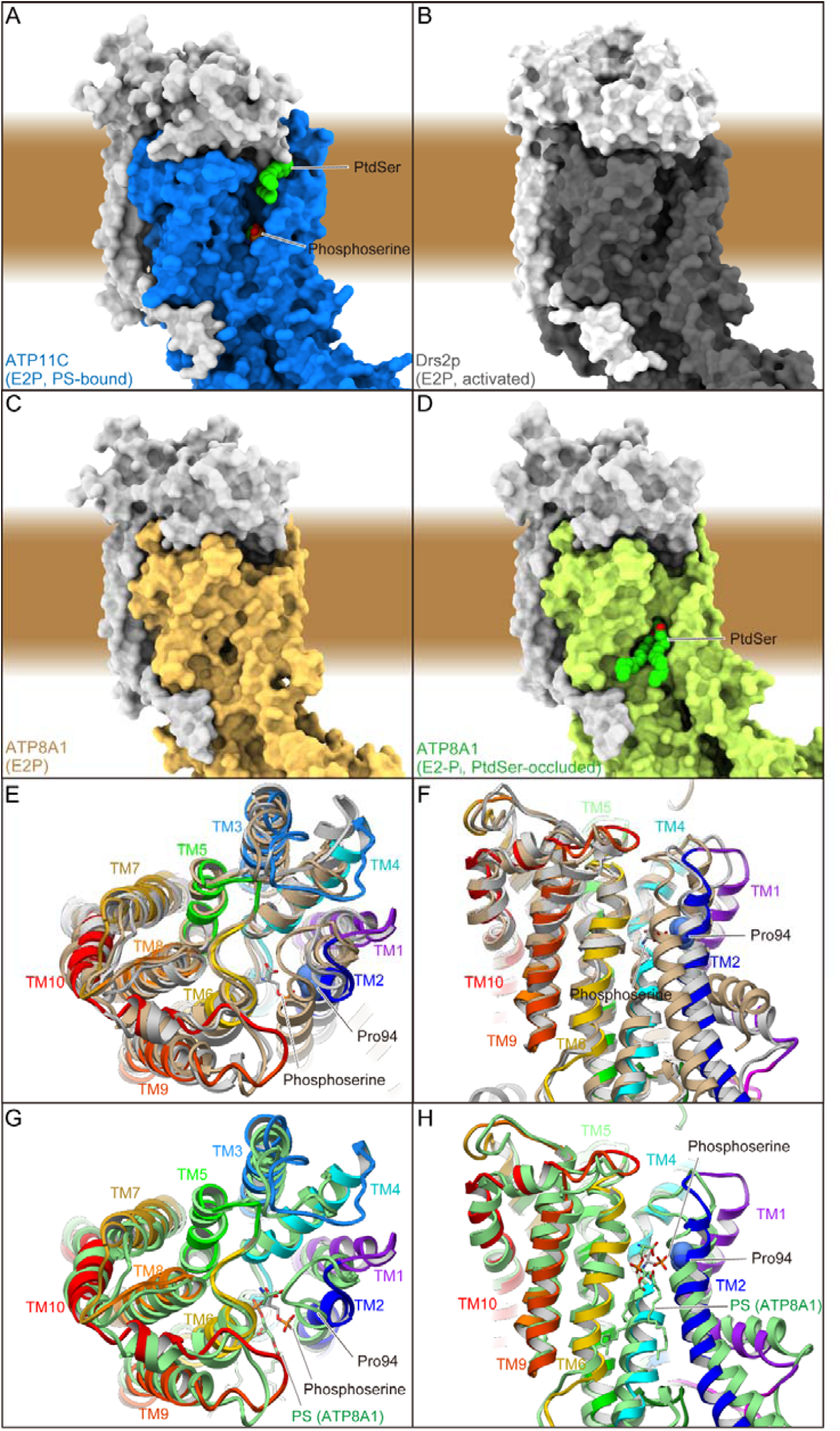
Comparison of membrane crevices. (A-D) Surface representation of the atomic models of ATP11C E2P state (A), Drs2p E2P activated form (B), ATP8A1 E2P (C) and E2-P_i_ state (D). PtdSer and phosphoserine are indicated as spheres. Brown background indicates approximate location of the lipid bilayer. (E-H) Comparison of the TM helix arrangement in ribbon representation. Atomic models of ATP11C (color codes as in Fig. 1), Drs2p (light grey) and ATP8A1 (tan) are aligned according to their TM helices (E,F). ATP8A1 E2-P_i_ transition state (light green) is also compared with ATP11C E2P state (G,H) Only catalytic subunits are shown in the figure, viewed from the exoplasmic side (E,G) or perpendicular to the membrane plane with exoplasmic side up (F,H). Phosphoserine (sticks) and Pro94 (spheres) in ATP11C, and PtdSer occluded in ATP8A1 E2-P_i_ state (sticks) are indicated in the figure.

**Fig. S7.**
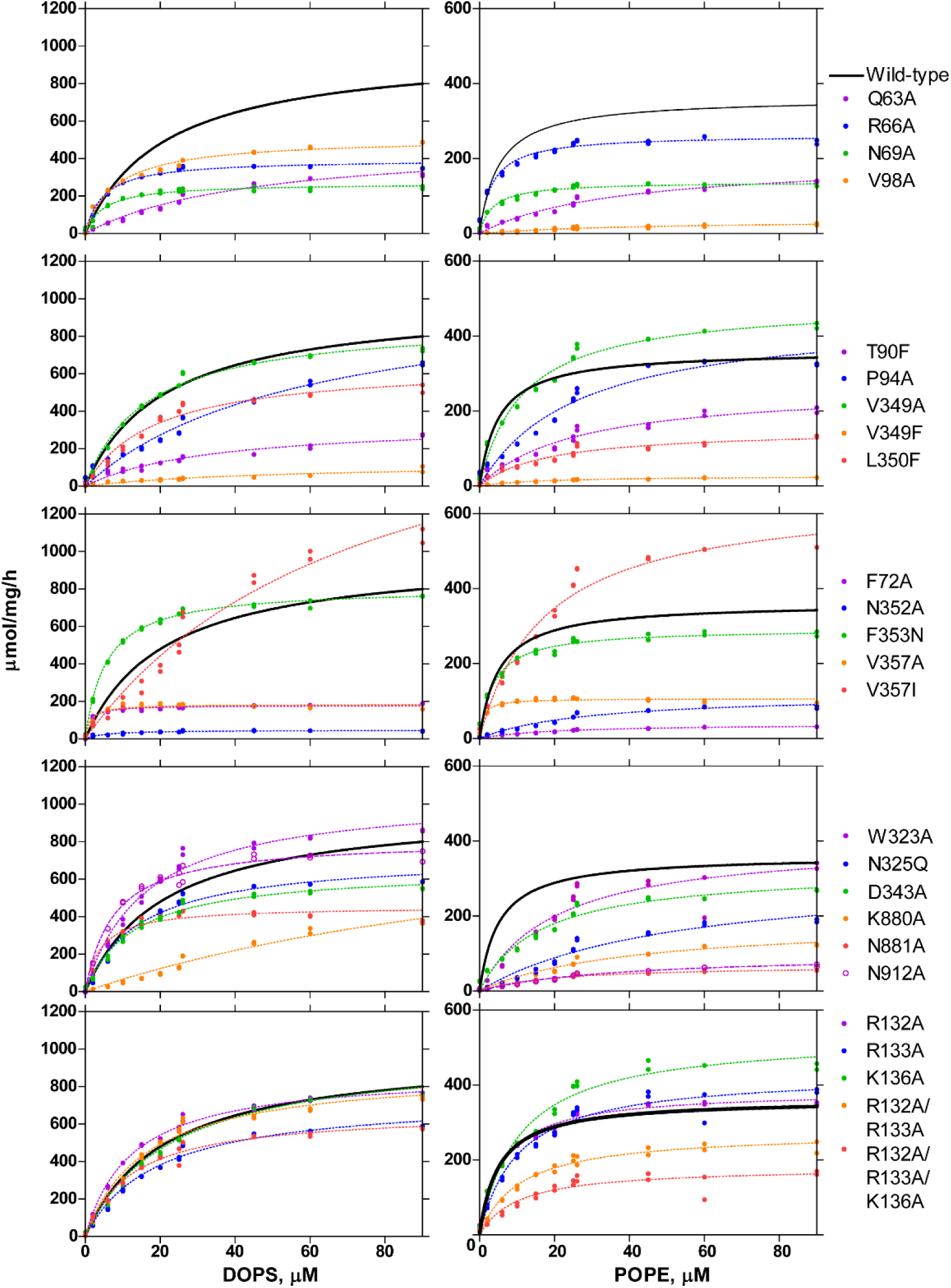
Phospholipid-dependence of ATPase activity for mutants. ATPase activities of indicated mutants are plotted as a function of DOPS (left) or POPE (right) concentration. Mutants are categorized as follows; cytoplasmic gate (1st row), surface of the membrane cleft (2nd row), occlusion site (3rd row), TM3-4 loop at exoplasmic cavity and residues in TM5 and 6 (4th row), and CDC50A exoplasmic domain facing the cavity (5th row). ATPase activity for the wild-type enzyme is shown in all graphs as a control (black lines).

**Table S1.**
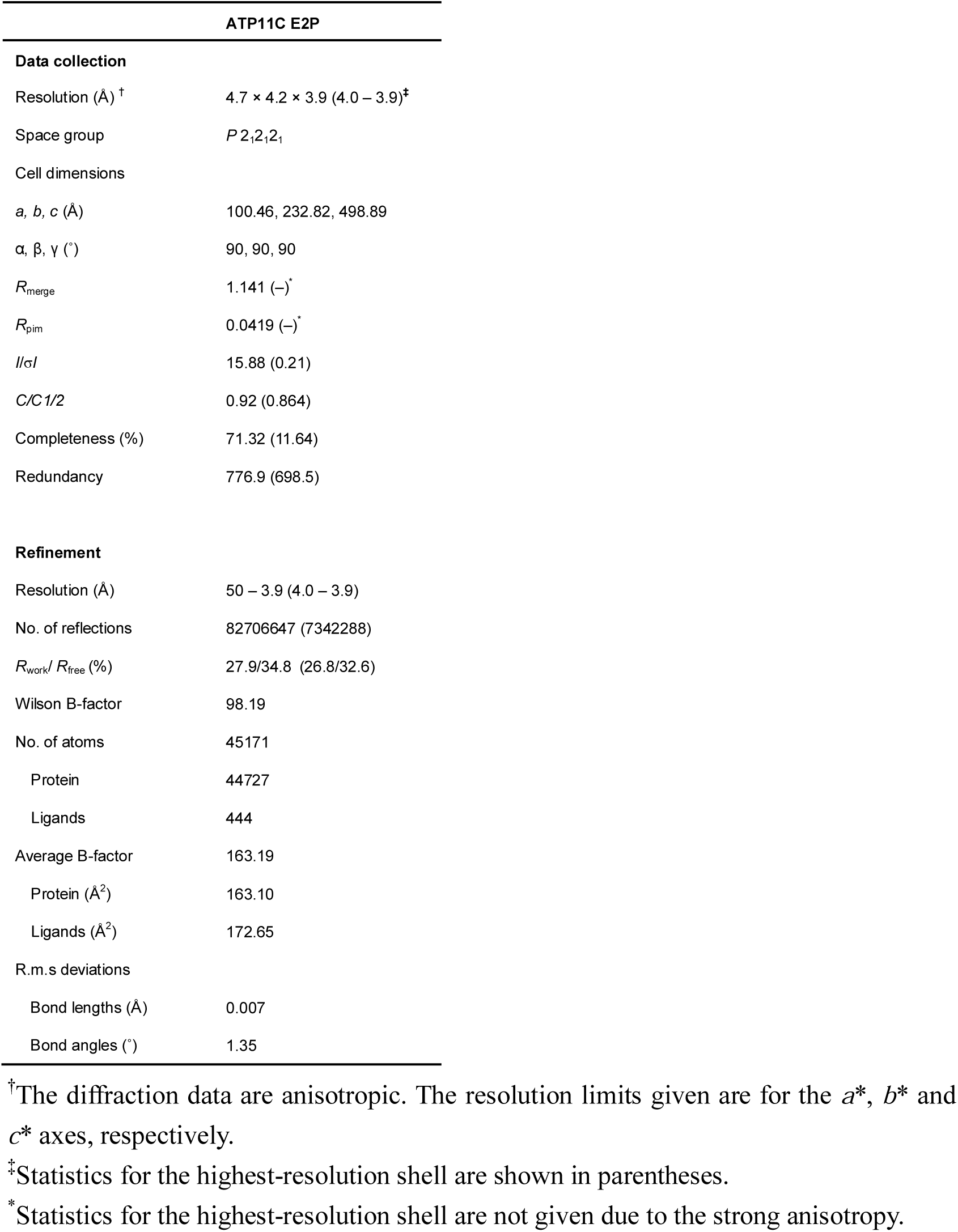
Data collection and refinement statistics.

